# Discovering triple negative breast cancers (TNBC) transcriptomic profiles via massive screening of single cell RNA-sequencing (scRNA-seq) datasets

**DOI:** 10.1101/2022.07.19.500610

**Authors:** Mohd Rifqi Rafsanjani, Muhammad Asyraf Onn, Norashikin Zakaria

## Abstract

Triple negative breast cancers (TNBC) account for 18% of the breast cancer patients in Malaysia and also account for one of the worst prognoses albeit it is the rare cancer types. TNBC is defined as breast cancers that tested negative on all 3 tests of oestrogen/progesterone (ER/PR) receptor and human epidermal growth receptor 2 (HER2). Understanding the TNBC biology is challenging due to the fact of multiple subtypes and its heterogeneity in patient samples. Here we attempt to examine the transcriptional profiling of TNBC using the single cell RNA-sequencing (scRNA-seq) analysis of different publicly available TNBC datasets. In comparison to bulk-RNA, scRNA-seq allowed the mapping of transcriptional profiling of TNBC at the single cell resolution and able to accurately pinpoint pan-transcriptional markers of TNBC.

## Introduction

The prevalence of the breast cancer among woman is excruciatingly high as it is the number one cancer for woman and Triple Negative breast Cancer (TNBC) is one of the breast cancer subsets that is difficult to treat due to its complexity. Study conducted by Malaysia cancer centre, demonstrate that 25 out of 143 breast cancer patients were Triple Negative Breast Cancer (TNBC) (Ahmad et al. 2019) indicating that TNBC among Malaysian population is at the level required attention. In metastasis state, it would be the worst survival outcome for the TNBC patients based on the multivariate analysis. TNBC is generally the breast cancers that tested negative on all 3 tests of oestrogen/progesterone (ER/PR) receptor and human epidermal growth receptor 2 (HER2). Despite being the rarest form of cancer overall, TNBC are more common in women below 40 years old, either black woman or with BRCA1 mutation (Abdul Aziz, Md Salleh, and Ankathil 2020). It is also more prevalent among demographics below 50 years old and with familial genetics. Unlike other types of invasive breast cancer such as adenocystic carcinoma, medullary carcinoma and other carcinomas, TNBC is more aggressive as it tends to enlarge and metastasize more rapidly with poor prognosis and fewer treatment options (Abdul Aziz et al. 2020).

Stratifying patients with TNBCs has only started since the past 10 years. Clustering of the gene expression data into six subtypes showing different responses into treatment but unfortunately there is no significant difference in the relapse-free survival statistically (Burstein et al. 2015; Lehmann et al. 2011; Yu et al. 2020). Metastatic cancer is also a persistent challenge in the treatment of breast cancer and its biological behaviours can be thoroughly understood by the application of recent development method of sequencing technology, the single-cell RNA sequencing (scRNA-seq) (Ren et al. 2021). scRNA-seq is one of the latest sequencing technologies at single cell resolution in comparison to bulk RNA-seq that sequence the cells at the tissue or ‘bulk’ of cells.

Mapping TNBCs to a single-cell resolution is vital as tumour growth is dependent on the diversity of cancer cell population and its microenvironment. Heterogenous long non-coding RNA (lncRNA) expression in xenografted TNBCs were accomplished with the use of 10x Genomics droplet-based technology subjected to its single-cell RNA sequencing. The transcriptomic data was analysed to obtain the inferred copy number aberration profiles via single-cell Copy Number Aberrations (scCNA). Further understanding on intratumor heterogeneity (ITH) which is a distinct cell population which are phenotypically and molecularly different is imperative in understanding key genetic vulnerabilities could be achieve via scRNA-seq (Pinkney, Black, and Diermeier 2021).

Here, we attempted to examine the transcriptomic profiling of TNBC from multiple scRNA-seq datasets where gene expression level could be observed at single cell resolution. At this high resolution possible in cellular biology, we wanted to understand the biological and cellular process happening inside the TNBC cells where perhaps we could pinpoint certain potential pathways to target. To do so, we pooled different publicly available sc-RNAseq from GEO database and further processed it using python-based single cell tool, Scanpy.

## Results and Discussion

### Overview of the datasets

From the 6 analysed TNBC scRNA-seq datasets, 5 were chosen for further analysis consisting of 3 cancer patients and 2 cell lines in separate analysis. GSE188600 is excluded from analysis due to lack of epithelial cells detected using our pipeline. Plus, the original paper was not focusing on the TNBC or epithelial cells, but more into immune cells study of TNBC cancer patients. Initial observation into these datasets shows additional cell types consisting of immune cells such as macrophages, B and T cells and additionally endothelial and stromal cells (Refer Figure 1.0). Leiden clustering demonstrate the different profile of the gene expressions and probably also the heterogeneity of the epithelial tissues that would be investigate later. The cells that did not satisfy any of the cell gene markers provided is labelled as unknown and would not be investigate further. Future unknown cells will also be excluded from the analysis. The epithelial cells detected in this fast-screening step consist of more than 50% of each GSEs (except the excluded GSE188600 which is less than 10%) and at best more than 90% in GSE143423 in patient samples, while in cell lines samples consists as high as 100% epithelial cells. Our initial screening provides easy and scalable screening of multiple TNBC datasets that allows us to screen a big number of datasets.

**Figure 1.**
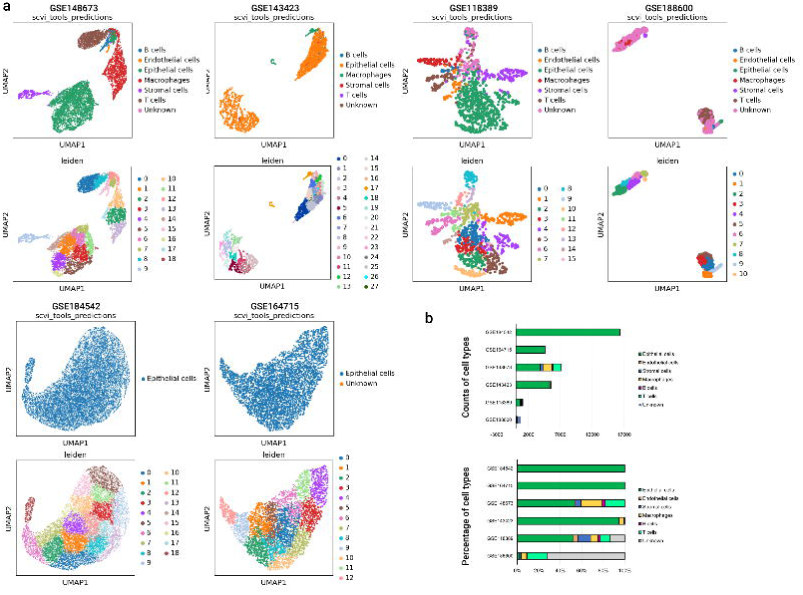
Screening of the different TNBC dataset from NCBI GEO. **a**) Cell types detected and number of clusters detected in each dataset where samples GSE188600, GSE118389, GSE143423 and GSE148673 is from TNBC patient samples and two others are from TNBC cell lines (GSE164715, GSE184542) hence only epithelial cells were detected. CellAssign tool was used for cell types annotation and Scanpy tools for data processing and graph plotting functions. **b**) Overview of each datasets summary showing the cell types count (top) and its percentage within its individual dataset (bottom).

These datasets than split into two scRNA-seq datasets of TNBC patients and TNBC cell lines that analysed separately. For the TNBC patients sample datasets, epithelial cells consist of more than 50% of overall cellular populations annotated using CellAssign were selected (Figure 2). Our clustering parameter were tune for high cell clustering as shown by the cluster number (in total 42 clusters). Since we are combining of huge data from multiple experiments and patient sample heterogeneity, we reasoned that high resolution clustering allows sensitive detection of annotated cell types or cell clusters that share similar transcriptome profiles. The annotated epithelial cells later split into its own dataset and combined with normal breast epithelial single cell dataset as a reference.

**Figure 2.**
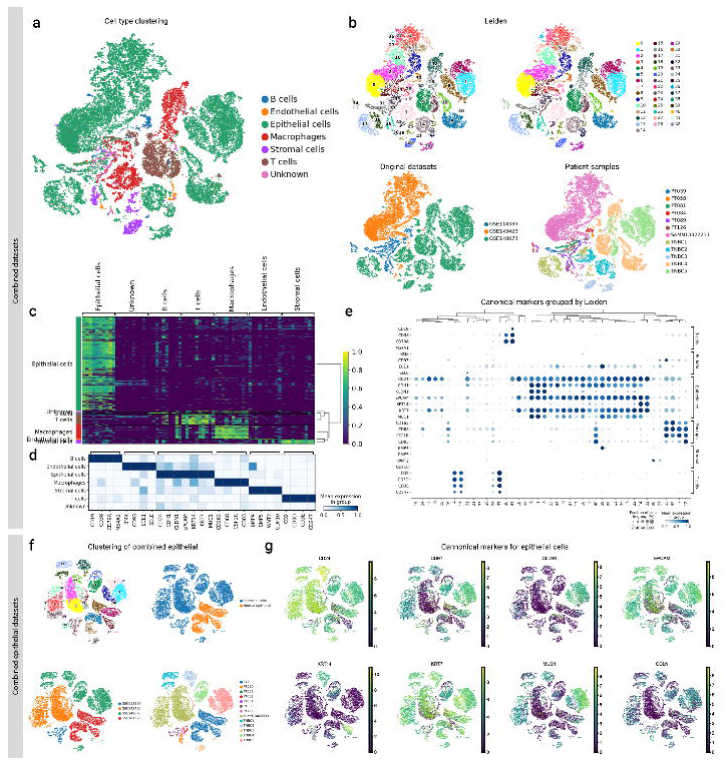
Combined dataset of TNBC scRNA-seq and epithelial cell selection. **a, b**) The cell clusters of combined datasets showing the notated cell types consisting of immune cells, stromal cells, endothelial and epithelial cells. Undetected cell types are marked as unknown cell types. UMAP plots also shown the cell clusters original GSEs as batch and individual patients. **c**) The ranked genes of the combined dataset as shown in heatmap showing the clustering of gene expression. **d**) The canonical marker genes of each cell types to validate the annotated cell types. **e**) Similar with **d** but with as shown as dot plot and cluster numbers showing relative mean gene expression and portion of those cells’ expression those genes. **f**) Epithelial cell dataset only subset from the overall dataset combined with normal breast epithelial cells. This dataset is then run for CNVs using InferCNVpy tool and CNV scores indicates those cells with high CNVs. These scores then will distinguish between normal epithelial and tumour epithelial cells and annotated as normal’ or ‘tumor’. Cycling cells also detected by computing the cell cycle scores based on the expressed genes. Tumour epithelial cell clusters contained all the phase indicating its cycling state while normal epithelial cell cluster show no or minimal G2M and S phase. **g**) Epithelial markers gene expression visualised via UMAP plots of the cell cluster.2

**Figure 3.**
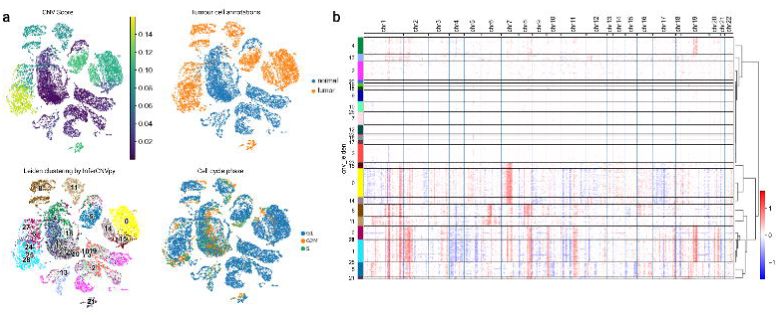
The copy number variation (CNV) and identification of cycling cells. **a)** Cell clusters indicating the CNV scores, annotated tumour cells, re-clustering based on the CNV scores and the cell cycle phase within the cell’s population. The CNV scores correlate to the origins of the TNBC patient sample. The cell cycles phase of the cycling cells are also correlates to the annotated tumour with exception. **b**) The heatmap plot visualising the cell clusters demonstrated the gain (red colour) and the lost (blue colour) of chromosomal number indicating the accurate annotation of both tumour and cell types. White indicates no difference to the normalepithelial breast cancer cells.

### Cell types annotations

Using CellAssign tool, we visualised the tumour microenvironment of TNBC samples data. Cannonical marker expression validate both annotated cells and its corresponding Leiden clustering as shown in the dot plots of Figure 2e. Additionally, these cells shared similar gene expression profiles as seen in the heatmap gene expression clusters. Since we want to focus on the epithelial cells, we also visualised the expression of epithelial canonical marker cell via UMAP plots as seen in Figure 2h. The epithelial clusters highly expressed CD24, KRT17 and EPCAM while some other clusters are selectively express KRT14, MUC1 and CLDN1. At this point, our highly confident of the CellAssign results annotating of these cell types and the epithelial cell types then isolated, combined with normal epithelial cell for the new combined epithelial datasets. This is necessary as normal breast epithelial cells will be used as reference for later analysis and to distinguish the gene expression profile between the TNBC and normal cells.

### TNBC tumour identification

To identify tumour cell within epithelial cells populations, we employ two step approach: 1) identification of CNV in comparison to the normal cells and 2) clusters containing cycling cells. CNV of chromosomes is one of the properties of cancerous cells. Using InferCNVpy tool, first the genes in the dataset is annotated into the chromosome position and the gene expression of the TNBC patient samples compared between the known normal epithelial gene expression. Each cell is given a CNV score (Figure 2) that indicates either gain or loss chromosomal copy number, the high the CNV scores the high probability of the cell gaining CNV. To accurately annotate the tumour cells, we re-clustering these cell populations using these scores and group into normal or tumour cells which is more reliable quantitative method rather than just using visualisation. Heatmap visualising these two clusters demonstrated the gain (red colour) and the lost (blue colour) of chromosomal number indicating the accurate annotation of both tumour and cell types. Visual validation also demonstrates that the cluster of epithelial cells from the normal patients has almost none of the CNV scores.

Additionally, we compared and validate our CNV analysis to the presence of cycling cells. Using Scanpy tool, we run ‘score gene cell cycle’ functions to annotate the cell phase of each cell either G1, 2M or S. As expected, all the tumour clusters have these cycling cells with all three cell cycle phase presence while the epithelial cluster from normal patients has almost none. This further validate our tumour cells annotations. On top of that, we also observed the presence of cycling cells within epithelial clusters that are not cancerous originate from the cancer patients. Since these cells has low CNVs as indicated by the CNV scores done previously, we conclude theses normal cycling epithelial cells are not as tumour.

## Methodology

### Dataset screening and compilations

scRNA-seq of TNBC datasets were obtained from NCBI Geo Omnibus Expression (GEO) and screened for epithelial cells counts. At least four TNBC datasets from patient samples (GSE188600, GSE118389, GSE143423, GSE148673) and two TNBC cell lines (GSE164715, GSE184542) were obtained for pre-screening using. Each data was annotated with its GSE numbers and patient’s or cell lines labelling using AnnData format and Scanpy tool, with minimal pre-processing step done to exclude low cells than contain less than 250 genes, not more than 20% mitochondrial gene content and only to include genes will counts less than 10,000. For cell types annotation, Cell Assign tool was used and the UMAP of these cell types were plotted as shown in Figure 1.0. Total of each cell types detected were counted where the epithelial cells count is less than 100 cells, it would be excluded for further analysis. We combined these selected datasets into two; one for the patient samples analysis and the other one is for cell lines analysis. Overall, we analysed 5 TNBC samples including 3 from patient samples and 2 from cell lines. For normal breast epithelial datasets that would be use for comparison in later step, we sampled scRNA-seq data from 3 patients (labelled as N1105, N280 and N-MH0064) from GSE161529 combined into one.

### Cell types annotation and validation

Throughout our analysis, we used CellAssign tool to annotate cell types in each dataset. CellAssign is a cell types automated annotation tool that use a probabilistic based method defined by set of marker genes. Hence, we used cell marker of tumour microenvironment based on the previous TNBC study (Wu et al. 2021). No normalisation or batch correction is done prior to CellAssign step since it required a raw count and will automatically adjust the any correction accordingly to the data. After annotation, principal component analysis (PCA), neighbours and clustering were computed using Scanpy prior the cell type UMAP were plotted. To validate our cell type annotation, we use a list of canonical marker expressions from PangoDB corresponding to each cell types and select at least four or more cell markers to verify CellAssign annotations. The canonical gene markers then were plotted into matrix and dot plots as seen Figure 2.

### Epithelial cell selection

Since our main objective is to obtain epithelial cells for downstream analysis, we additionally compare and check the canonical gene markers of epithelial cells to the Leiden clustering to identify correct epithelial gene expressions. The annotated epithelial cells type was split into its own dataset and combined with normal breast epithelial scRNA-seq dataset prepared in the first step. This combined epithelial scRNA-seq data then undergoes re-clustering using scVI-tools, the same computational tools used by CellAssign, but without additional gene marker annotations. This will allow similar gene expression profiles within epithelial cells to cluster together based on the scVI-tools probabilistic modelling which is a crucial component in our analysis since we are analysing diverse TNBC sample origins with mixed tumour and non-tumour cell types within.

### Cell cycle analysis

Cell cycle analysis were done to identify cycling cells within their epithelial populations. A gene set indicating gene cycle phase as characterized by (Tirosh et al. 2016) were used and scored against using score gene cell cycle functions in Scanpy. As the results, each cell was scored against a cell cycle phase whether G1, G2M or S. Epithelial clusters that has high G2M and S scores is considered as a cycling cell while non cycling epithelial cluster is considered only those with G1 scores.

### Copy number variations (CNVs)

To classify which epithelial cells is a tumour, we performed CNV analysis using Python based tool, InferCNVpy. The genes of combined epithelial scRNA-seq dataset earlier were annotate against human chromosome GRCh38.p13 available at GENECODE (https://www.gencodegenes.org/human/) where comprehensive gene annotation of GTF that cover CHR regions was used. Then the CNVs analysis were done with normal epithelial cells as the reference. Heatmap of CNVs within chromosomes were also plotted to visualise either gain or loss of chromosomal copy number. Finally, tumour cells were classified based on the CNV score and CNV’s clustering before annotated either as tumour or normal cells.

## Acknowledgements

The authors would like to express their immense gratitude to Ahmad Mustakim Mohamad, School of Distance Education, Universiti Sains Malaysia for their statistical support.

## Notes

### Competing Interest Statement

The authors have declared no competing interest.

